# Self-amplifying ROS-sensitive SN38 Dimeric Prodrug nanoparticles for Combined Chemotherapy and Ferroptosis in Cancer Treatment

**DOI:** 10.1101/2024.01.22.576778

**Authors:** Yu Qin, Fenghui Wang, Zeping Gao, Chutong Tian, Ken-ichiro Kamei

**Author notes:** Corresponding authors: E-mail addresses (C. T.); (K. K.).

## Abstract

Chemotherapy drugs are often limited by their own clinical shortcomings and serious adverse consequences. To solve these problems, we developed a self-amplifying reactive oxygen species (ROS)-sensitive dimeric prodrug nanoparticles, namely SN38-CA@FC NPs for tumor treatment. A ROS-sensitive 7-ethyl-10-hydroxycamptothecin (SN38) prodrug (SN38-CA) was synthesized by a thioacetal linker between SN38 and the ROS generator cinnamaldehyde (CA). The subsequent release of SN38 inflicts DNA damage, exerting chemotherapeutic effects, while the liberated CA intensifies ferroptosis through Fenton reaction-mediated disruption of the redox balance. This dual-action strategy not only leverages chemotherapy but also induces ferroptosis, establishing a synergistic therapeutic paradigm. The system is uniquely characterized by a positive feedback loop where ROS instigates the release of SN38/CA, which in turn promotes further ROS production. In experimental evaluations, this combination therapy exhibited potent antitumor activity against both A549 and LLC cancer cell lines, as well as in xenograft LLC-bearing C57BL/6 mouse models. Collectively, our findings introduce a transformative Nano-Drug Delivery System (NDDS) that holds significant promise for advancing cancer chemotherapy and ferroptosis-based therapies.

## 1. Introduction

Chemotherapy remains a cornerstone in tumor treatment strategies, and Camptothecin (CPT) has risen to prominence for its proficiency in inhibiting DNA damage through topoisomerase 1 [1]. Among its derivatives, 7-Ethyl-10-hydroxycamptothecin (SN38) stands out, mirroring CPT’s mechanism and earning recognition as one of the most potent antitumor agents within this family [2]. SN38 has been reported to be 100-1000 times more effective against cancer *in vitro* than irinotecan (CPT-11) [3]. However, the clinical application of SN38 as an anticancer drug is significantly hindered by its poor aqueous solubility, inadequate stability and pronounced systemic toxicity [3].

Recent advancements have sought to mitigate these clinical challenges by innovating SN38 prodrugs and nanomedicine approach. The NanoDrug Delivery Systems (NDDS) for SN38, in particular, offer notable improvements in water solubility, bioavailability [4, 5], tumor targeting via enhanced permeability and retention (EPR) effect [6], and the potential of combination therapies [7]. Despite these advantages, hurdles persist, including suboptimal drug loading, inconsistent in vivo conversion rates, and variable toxicity profiles [5]. Furthermore, reliance on chemotherapy alone can result in tumor resistance, recurrence, and metastasis, burdening patients with severe side effects post-treatment [8].

In response, there is an emerging focus on designing SN38 prodrugs with controlled release mechanisms and constructing nanoplatforms that integrate additional therapeutic modalities [9]. Enter ferroptosis—a distinct mode of cell death driven by iron-dependent lipid peroxide accumulation—offering a novel avenue for targeting chemotherapy-resistant tumors [10]. The Fenton reaction, a catalyst for ferroptosis, leverages ferrous ions (Fe^2+^) to convert hydrogen peroxide (H_2_O_2_) within the tumor microenvironment into cytotoxic hydroxyl radical (·OH), resulting in a large amount of LPOs accumulation. Despite the promise of ferroptosis induction via the Fenton reaction, its efficacy is contingent on intratumoral hydrogen peroxide levels [11] and can be compromised by intracellular glutathione (GSH) [12].

To bolster this ferroptotic pathway, we turn to Cinnamaldehyde (CA), derived from traditional cinnamon bark oil, known for its ability to deplete GSH and augment hydrogen peroxide in tumor cells [13–15]. CA facilitates this by engaging in a Michael-type addition (1,4-addition) with GSH to the double bonds at α, β-position of the aldehydic functionality of CA, resulting in the conversion of GSH into oxidized glutathione (GSSG) [16, 17] and enhancing the oxidative stress within the tumor [18].

Here we introduce ROS-sensitive SN38 dimeric prodrug nanoparticles, namely SN38-CA@FC NPs, harnessing the dual potency of chemotherapy and ferroptosis (Fig. 1). Through a novel synthesis of SN38-CA coupled with Ferrocene carboxaldehyde (FC), we formulate uniform nanoparticles designed to target tumors effectively [19]. These nanoparticles exploit the EPR effect for tumor specificity [20] and leverage the tumor’s ROS-rich environment to release SN38 and CA. This release initiates a dual-action: SN38’s chemotherapeutic inhibition of topoisomerase 1 and CA’s amplification of the Fenton reaction, culminating in heightened ferroptosis. The resulting hydroxyl radicals, in a self-amplifying loop, further promote the decomposition of SN38-CA, embodying a positive feedback mechanism of ROS-triggered drug release and drug-mediated ROS generation —a strategy we believe could substantially elevate the efficacy of tumor treatments. Our comprehensive evaluation of SN38-CA@FC NPs spans both 2D cellular models and in vivo animal studies, aiming to demonstrate the profound potential of this innovative therapeutic approach. This integration of chemotherapy and ferroptosis via a nanoparticle delivery system could represent a significant leap forward in the ongoing battle against cancer.

**Fig. 1.**
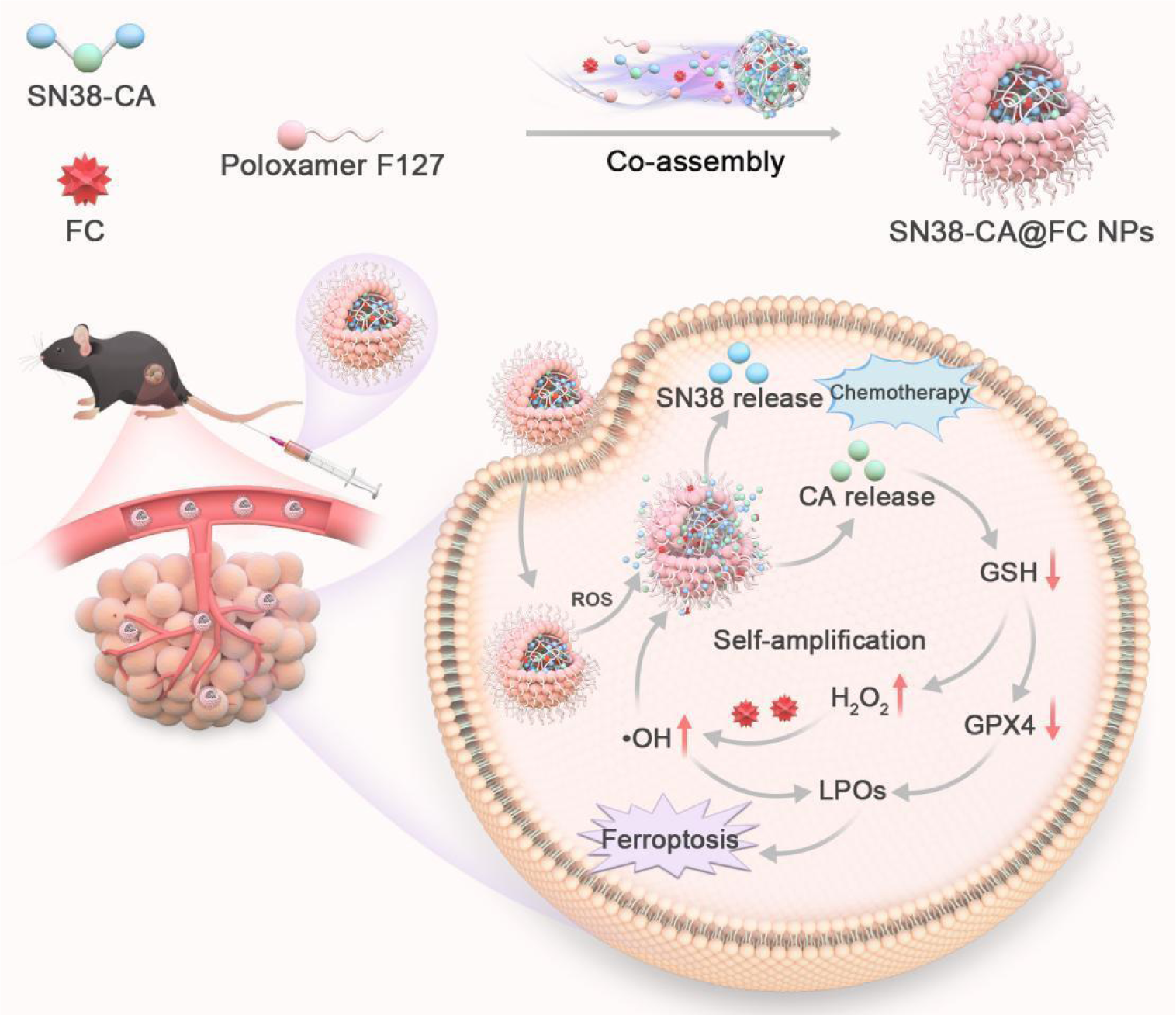
Schematic diagram of the self-assembly process of the SN38-CA@FC NPs and its proposed mechanism for combination of chemotherapy and ferroptosis.

## 2. Results and discussion

### 2.1 Chemical synthesis

To solve the clinical defects of SN38 and simultaneously amplify the efficiency of the Fenton reaction, we conjugated SN38 and CA into ROS-sensitive prodrug, denotated as SN38-CA, using a thioacetal linker (Fig. 2). The synthetic pathway commences with the First, the aldehyde group of CA is replaced by two hydroxyl groups with 2-Mercaptoethanol, and then the resulting midbody 1 is esterified with 4-Nitrophenyl chloroformate. Finally, SN38 could be coupled to CA through esterification with midbody 2. According to ^1^H-NMR and Mass spectrum, SN38-CA was successfully synthesized (Fig. S1-4). The SN38-CA structure consists of two SN38 and is a dimer structure, which greatly improves the drug loading of the nanoparticles [21].

**Fig. 2.**
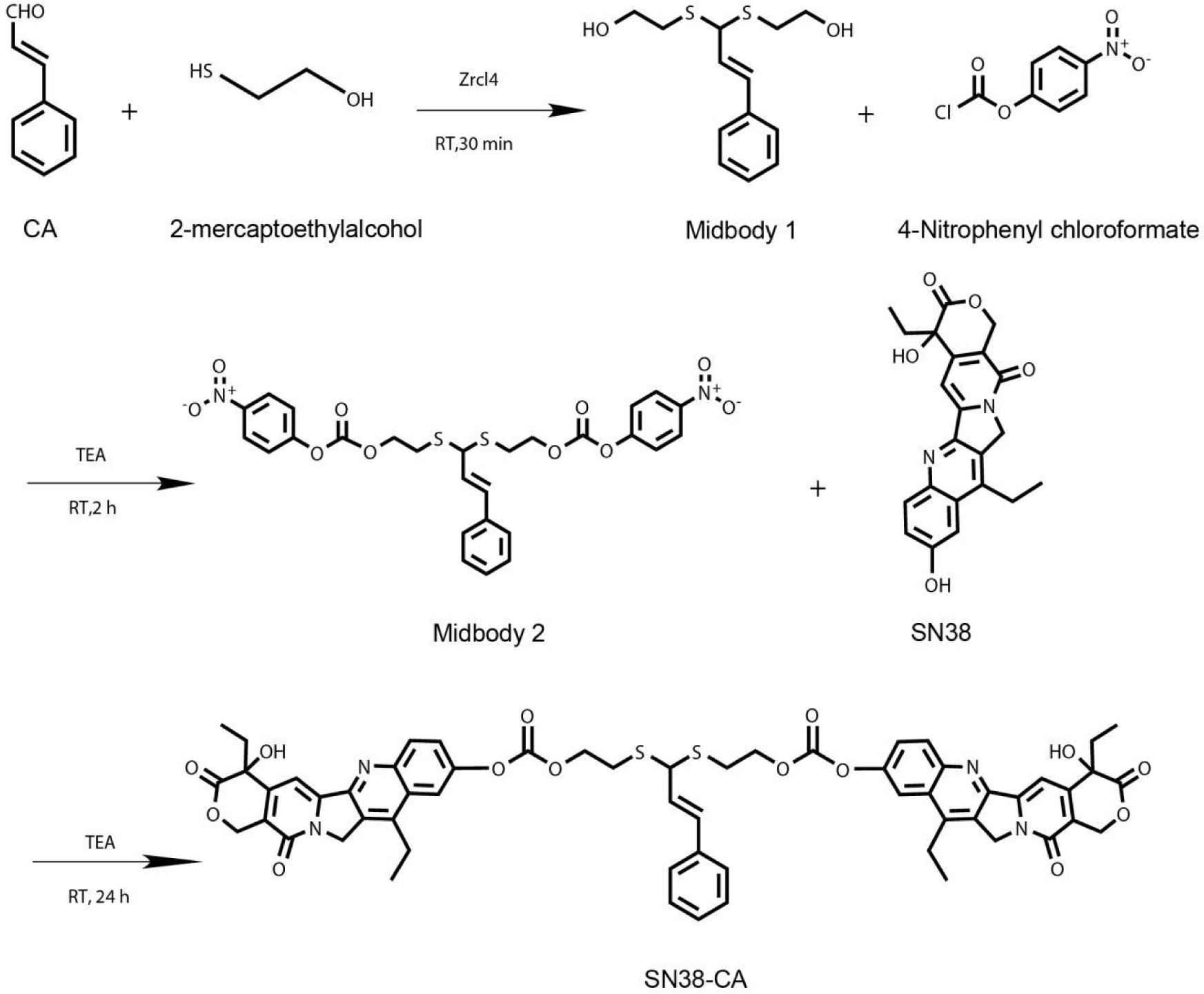
The synthetic route of SN38-CA.

The tumor microenvironment, characterized by elevated levels of reactive oxygen species (ROS), acts as a stimulus for the cleavage of SN38-CA, leading to the liberation of the parent drug. While the inherent ROS levels in tumors may not be abundant, the released CA functions as a generator of ROS, instigating a cascade that further propels the release of the prodrug. This establishes a positive feedback loop, a “trigger-release-trigger” mechanism. Integration with FC introduces highly active ·OH produced by the Fenton reaction, which intensify and expedite this feedback loop, optimizing the therapeutic strategy.

### 2.2 Preparation and Characterization of prodrug nanoparticles

To compare the anti-tumor effects of ferroptosis, we prepared two prodrug nanoparticles containing SN38-CA: SN38-CA NPs and SN38-CA@FC NPs. Nanoparticles were fabricated based on a one-step nanoprecipitation method[22], only an amount of Poloxamer F127 (20%, w/w) was used to increase the stability of the nanoparticles [23]. The particle sizes of SN38-CA NPs and SN38-CA@FC NPs are evenly distributed, ranging from 120.0 to 140.0 nm and 90.0 to 110 nm, and PDI ranges from 0.190 to 0.210 (Fig. 3 A-B). The zeta potential of SN38-CA NPs and SN38-CA@FC NPs were both negative, which was conducive to the uptake of nanoparticles by cells [24] (Fig. 3A-B). Images taken by TEM show that SN38-CA NPs and SN38-CA@FC NPs have uniform particle sizes and smooth surfaces (Fig. S5). Nanoparticles have colloidal properties, after red laser irradiation, both SN38-CA NPs and SN38-CA@FC NPs exhibited a significant Tyndall effect (Fig. S6). In order to verify the Fenton reaction, FC was added to the solution of crystal violet and H_2_O_2_. Subsequently, the absorbance of crystal violet decreased significantly, indicating that Fenton reaction can occur between FC and H_2_O_2_ (Fig. S7-8). SN38-CA NPs and SN38-CA@FC NPsexhibited stability in PBS with 10% serum in 12 h, indicating that the prodrug nanoparticles could maintain stability in the blood circulation *in vivo* (Fig. 3C).

**Fig. 3.**
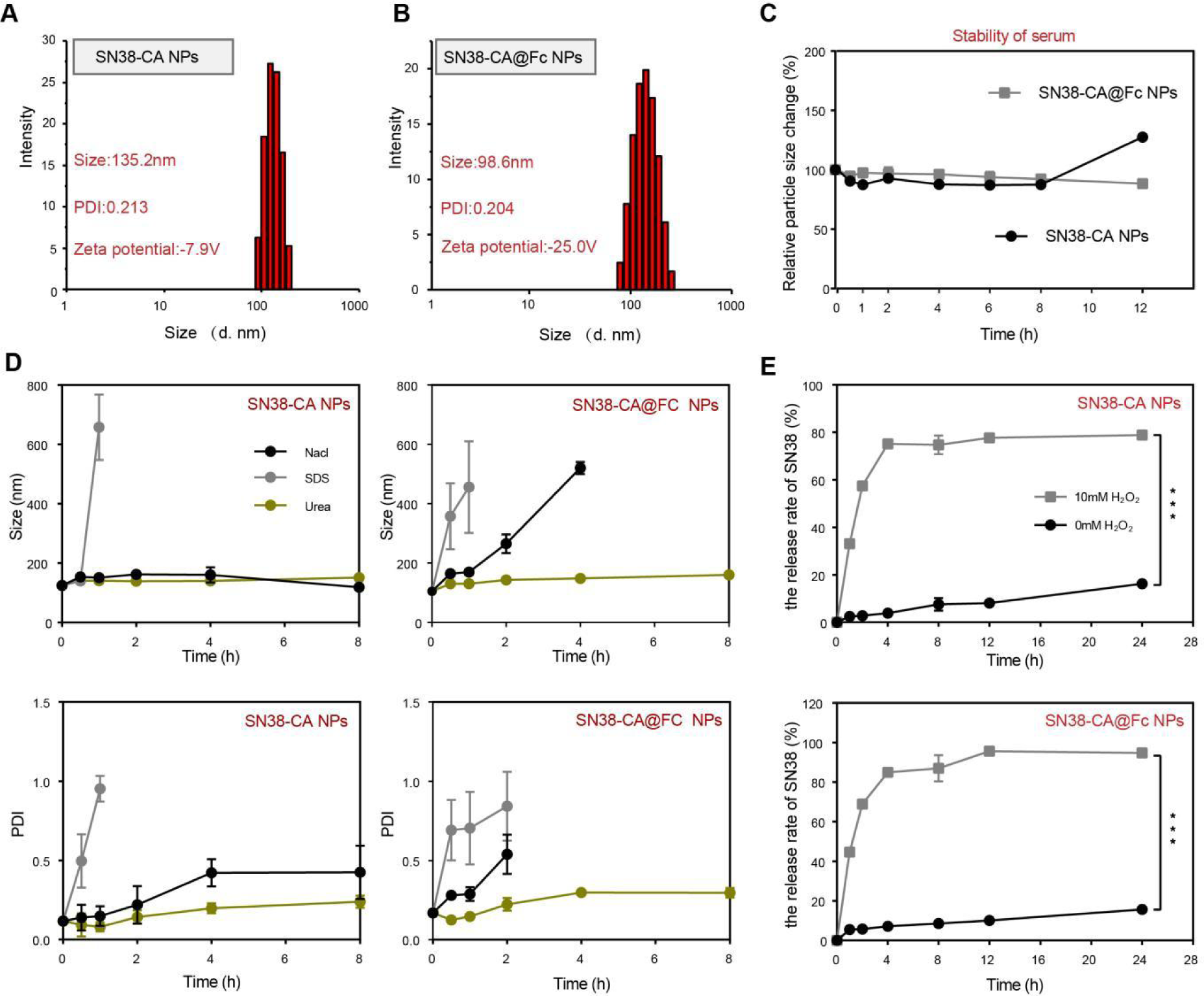
Characterization of SN38-CA NPs and SN38-CA@FC NPs. (A) Particle size distribution of SN38-CA NPs. (B)Particle size distribution of SN38-CA@FC NPs. (C) Stability of nanoparticles after incubation in PBS (pH 7.4) containing 10 % FBS at 37 °C. (D) Investigation of molecular interaction mechanisms of the assembly of nanoparticles. (E) Drug release rate of SN38 from SN38-CA NPs and SN38-CA@FC NPs under H_2_O_2_ stimulation. ns: no significance, ^∗^*p* < 0.05, ^∗∗^*p* < 0.01, and ^∗∗∗^*p* < 0.001.

### 2.3 Interactions Governing the Assembly of SN38-CA@FC Nanoparticles

Nanomedicine has revolutionized tumor therapy by offering alternatives to traditional nanocarrier-based drug delivery systems. Notably, carrier-free nanodrug delivery systems have garnered attention due to their high drug load, straightforward preparation process, and minimal toxicity. The assembly of these carrier-free systems is typically governed by non-covalent interactions, such as hydrophobic interactions, hydrogen bonding, and electrostatic forces [25].

To elucidate the specific interactions facilitating the assembly of SN38-CA@FC NPs, we challenged the molecular assembly with various salts designed to disrupt these forces. Our experiments utilized NaCl, SDS, and urea to target electrostatic, hydrophobic, and hydrogen bonds, respectively. According to the data presented in Fig. 3D, the particle stability remained unaltered in the presence of urea, suggesting minimal involvement of hydrogen bonding. In contrast, SDS addition resulted in increased particle size and polydispersity index (PDI) for both SN38-CA NPs and SN38-CA@FC NPs, indicating significant hydrophobic interactions. Similarly, the introduction of NaCl led to an increase in particle size and PDI for SN38-CA@FC NPs. While SN38-CA NPs did not exhibit a change in particle size, an increase in PDI was observed, implicating the influence of electrostatic interactions to a certain degree.

These findings imply that the assembly of SN38-CA@FC NPs is a concerted effort of hydrophobic and electrostatic interactions, with a dominant contribution from hydrophobic forces. This is likely because in an aqueous environment, hydrophobic components, such as aromatic rings, naturally tend to evade water and aggregate, which is a key driver for the nanoparticle assembly process. Moreover, the combination of covalent and non-covalent assembly methods observed in systems like SN38-CA@FC NPs is a common strategy. Here, multiple drugs are first linked via covalent bonds to form stimulus-responsive prodrugs, which are subsequently assembled into nanoparticles through non-covalent interactions. This dual-assembly approach underscores the intricate design and functional sophistication of current nanomedicine platforms [26].

### 2.4 ROS-Triggered Release Efficiency of SN38 from Nanoparticle Prodrugs

To evaluate the release efficiency of SN38 from SN38-CA NPs and SN38-CA@FC NPs in vitro triggered by ROS, we conducted in vitro assays using H_2_O_2_ as a ROS stimulus. The results, delineated in Fig. 3E, revealed a clear time-dependent release pattern of SN38 from both SN38-CA NPs and SN38-CA@FC NPs. Initially, upon H_2_O_2_ stimulation, there was a rapid liberation of SN38, which plateaued after 4 h. Following a 24-h incubation with 10 mM H_2_O_2_, SN38-CA NPs had released 78.88% of their SN38 content, while SN38-CA@FC NPs had released a notably higher percentage of 94.80%. In contrast, in the absence of H_2_O_2_, the release rates were considerably reduced; SN38-CA NPs released only 16.28% and SN38-CA@FC NPs released 15.71% of their SN38 after the same incubation period.

These observations confirm the robust ROS responsiveness of the SN38-CA formulation. The intrinsic self-amplifying mechanism of the nanoparticles ensures a rapid dissemination of the parent drug, SN38. Notably, SN38-CA@FC NPs demonstrated superior release metrics compared to SN38-CA NPs. This enhanced performance is likely due to ·OH produced by the Fenton reaction, which represent a more potent form of ROS, thereby amplifying and hastening the self-amplification process.

### 2.5 Cellular uptake and *in vitro* anti-tumor activity of SN38-@FC NPs

Recognizing the vital importance of cellular internalization for the anti-tumor efficacy of nanoparticles, we delved into assessing the cellular uptake efficiency of our nanoparticles. For this purpose, we utilized Coumarin-6 (C6)-labeled nanoparticles (SN38-CA@FC-C6 NPs), enabling the tracking of their internalization within the cells. A549 cells were subjected to treatment with both C6 Sol and SN38-CA@FC NPs, with subsequent monitoring of the uptake via fluorescence intensity measures. As depicted in Fig. 4A, both Ce6 and SN38-CA@FC NPs demonstrated a time-dependent cellular accumulation, with the fluorescence intensity at 4 h surpassing that at 1 h. Remarkably, SN38-CA@FC NPs exhibited a greater degree of accumulation in A549 cells than C6 Sol at both time points. This interaction between the cells and nanoparticles is influenced by nanoparticle physical attributes—size, shape, surface charge—and the nature of the cell membrane [27]. The optimized particle size distribution, appropriate size, and negative surface charge of SN38-CA@FC NPs are conducive to their efficient uptake.

**Fig. 4.**
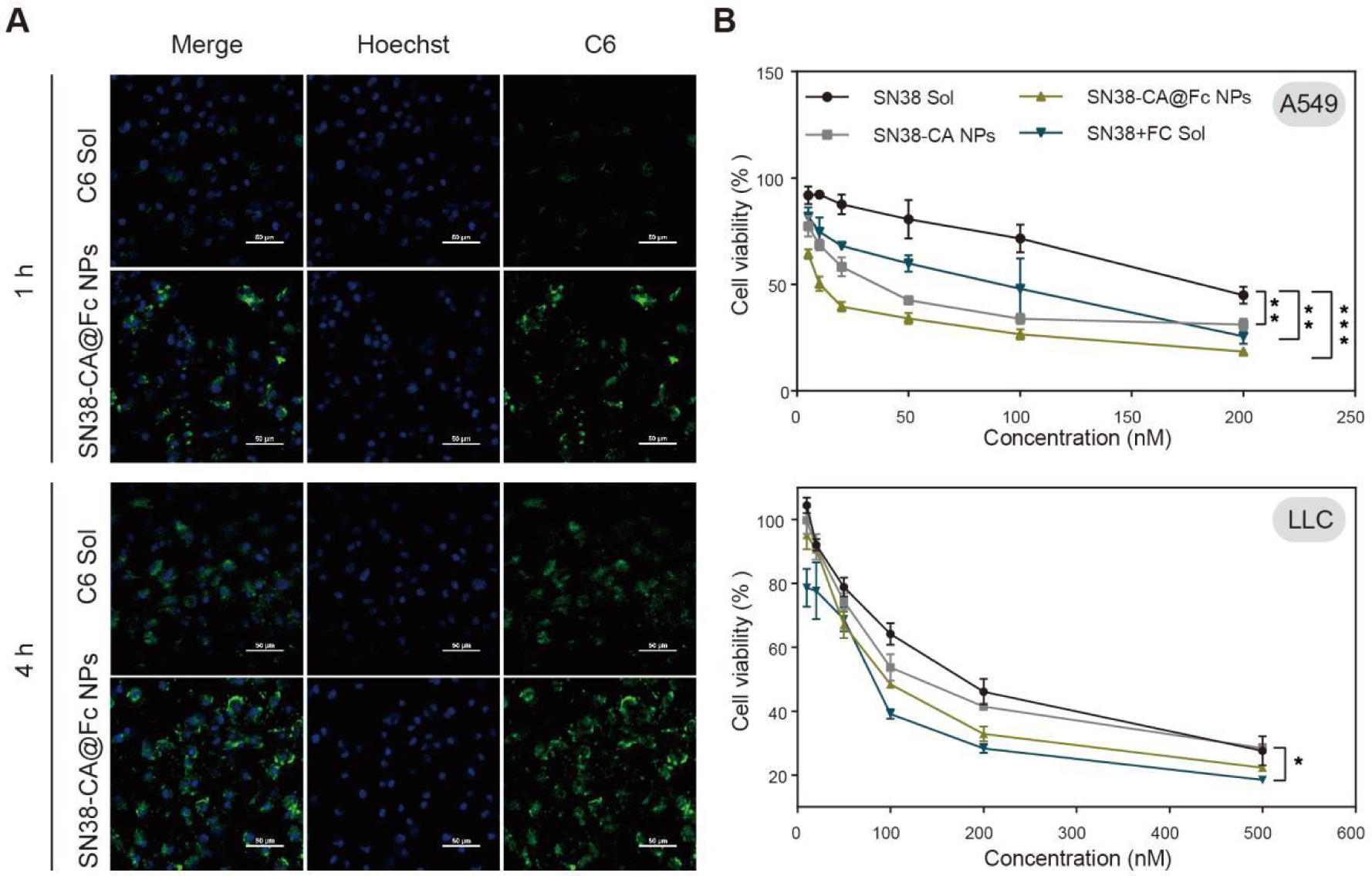
*In vitro* cellular level evaluation of nanoparticles. (A) Fluorescent micrographs to show the uptake capacity of C6 Sol and SN38-CA@FC-C6 NPs in A549 cells at 1 and 4 h; (B) Cell viability of SN38 Sol, SN38@Fc Sol, SN38-CA NPs, and SN38-CA@FC NPs against A549 and LLC cells at different concentrations after 48-h incubation. ns: no significance, ^∗^*p* < 0.05, ^∗∗^*p* < 0.01, and ^∗∗∗^*p* < 0.001.

Further, to probe the antitumor potential of SN38-CA@FC NPs *in vitro*, we assessed the cytotoxic effects against A549 and LLC tumor cells using the MTT assay. Fig. 4B demonstrates that all four drug formulations—SN38 Sol, SN38+Fc Sol, SN38-CA NPs, and SN38-CA@FC NPs—exhibited dose-dependent cytotoxicity across both cell lines. Notably, the toxicity patterns were consistent in A549 and LLC cells for all groups. SN38 Sol exhibited the least cytotoxicity, likely due to its known clinical shortcomings and suboptimal cellular uptake. In contrast, SN38+FC Sol displayed enhanced toxicity, which can be attributed to the Fenton reaction induced by FC, leading to increased cell death.

The nanoparticle formulations, SN38-CA NPs and SN38-CA@FC NPs, showed significantly higher cytotoxicity compared to the solution groups. This improved performance is credited to the nanoparticles’ superior cellular uptake and the prodrug modifications that enhance chemotherapy effectiveness. Among them, SN38-CA@FC NPs stood out, demonstrating exceptional cytotoxicity against both A549 and LLC tumor cells. This remarkable antitumor activity is the result of a synergistic effect combining chemotherapy with ferroptosis induction and a self-amplifying release mechanism, marking a significant advance in nanoparticle-mediated cancer therapy.

### 2.6 SN38-CA@FC NPs promote intracellular ROS accumulation

The intracellular release of CA from our nanoparticle system is engineered to deplete GSH and generate H_2_O_2_, a precursor for ·OH produced via the Fenton reaction. To assess the impact of this mechanism, A549 cells were treated with various formulations, and resultant ROS levels were monitored using the DCFH-DA fluorescent probe.

As shown in Fig. 5A-B, indicate that standalone FC induces negligible changes in ROS levels compared to the control PBS group. This is likely due to the limited Fenton reaction within the low-ROS cellular environment, resulting in minimal ·OH production. Conversely, SN38-CA NPs instigate a slight increase in ROS, even in the absence of FC. This suggests that CA can initiate a modest self-amplification of ROS, leading to an increase in both CA release and subsequent intracellular ROS levels.

**Fig. 5.**
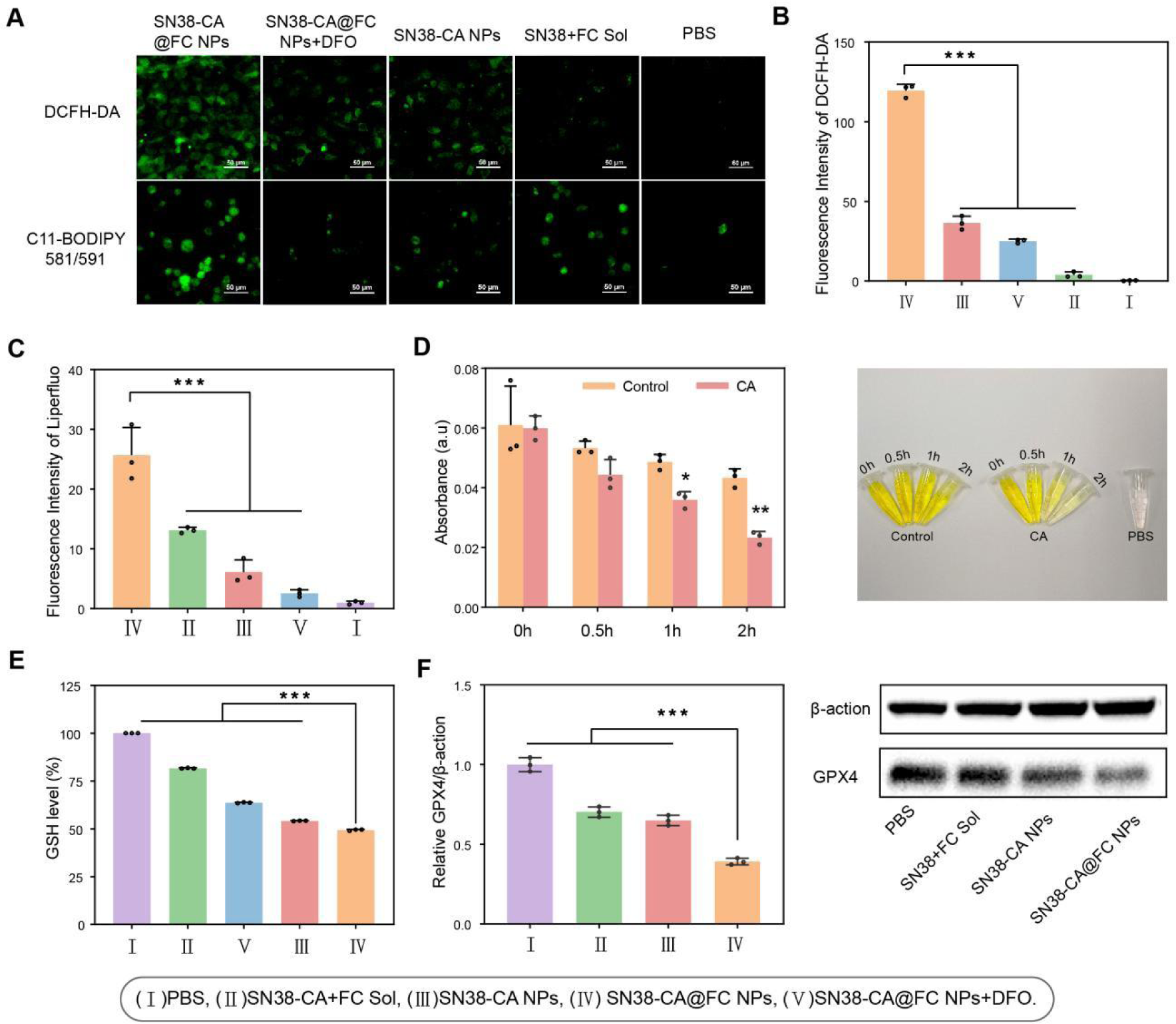
Impact of SN38-CA@FC NPs on Intracellular ROS, LPOs, GSH, and GPX4 Levels in A549 cells. (A) Fluorescence microscopy images illustrating the intracellular accumulation of LPOs in A549 cells treated with various samples, using C11-BODIPY ^581/591^ staining. Scale bars represent 50 μm. (B) Quantitative microscopic image analysis of intracellular ROS levels in A549 cells. (C) Quantitative microscopic image analysis of intracellular LPO levels in A549 cells. (D) Kinetic analysis of CA’s GSH depletion capability, as indicated by the time-dependent increase in DTNB-reactivity, suggesting effective GSH consumption by CA. (E) Bar graph showing the intracellular GSH levels in A549 cells following treatment with various samples. (F) Western blot analysis revealing the levels of GPX4 protein in A549 cells after treatment with different nanoparticle formulations. ns: no significance, ^∗^*p* < 0.05, ^∗∗^*p* < 0.01, and ^∗∗∗^*p* < 0.001.

The most substantial rise in ROS was seen with SN38-CA@FC NPs, underscoring the synergistic effect of the released CA in conjunction with FC. This combination fosters a broad self-amplifying cycle, markedly enhancing the release of CA, optimizing the efficiency of the Fenton reaction, and culminating in significant ROS production. The impact of the Fenton reaction was further confirmed by the introduction of the iron chelating agent, Deferoxamine (DFO), to the SN38-CA@FC NP treatment. Following the addition of DFO, a noticeable reduction in ROS levels was observed.

In summary, the data robustly demonstrate that SN38-CA@FC NPs can profoundly escalate intracellular ROS levels, indicative of a substantial disruption of the cellular redox system. This redox imbalance is a strategic target in our nanomedicine design, intended to amplify oxidative stress within tumor cells and thereby enhance anticancer efficacy.

### 2.7 SN38-CA@FC NPs promote Intracellular accumulation of LPOs

Ferroptosis is hallmarked by the accretion of lipid peroxides (LPOs), and to evaluate this in the context of our nanoparticles, A549 cells were treated with various samples. We employed the C11-BODIPY ^581/591^ probe to quantify intracellular LPO levels. According to Fig. 5A, B and C, SN38-CA@FC NPs exhibited the highest fluorescence intensity, signifying the most substantial LPO accumulation and consequent induction of ferroptosis in tumor cells. The addition of the iron chelator Deferoxamine (DFO) to the SN38-CA@FC NP treatment resulted in a notable decrease in LPO accumulation, underscoring the iron-dependence of this process. It was observed that the SN38+FC Sol group also facilitated LPO accumulation, albeit to a lesser extent. Interestingly, SN38-CA NPs, even without FC, led to a minor increase in LPOs, likely due to the indirect effects of CA, which disrupts GSH and GPX4 regulation. In summary, SN38-CA@FC NPs are capable of inducing significant LPO accumulation in tumor cells, thereby driving the cells towards ferroptosis.

### 2.8 SN38-CA@FC NPs induce intracellular GSH depletion

GSH reactivity, as evidenced by its reaction with 5,5’-Dithiobis(2-nitrobenzoic acid) (DTNB) to form a yellow compound, served as a preliminary test for CA’s GSH depletion capability, which exhibited a time-dependent consumption profile. To assess the impact of our nanoparticles on intracellular GSH levels, A549 cells were incubated with different treatments, and GSH concentrations were measured using a GSH assay kit. The data presented in Fig. 5E revealed a decrease in GSH levels across all groups compared to the control PBS group. Notably, the SN38-CA@FC group showed the most pronounced decline in GSH, pointing to the potent effect of ·OH generated by the Fenton reaction, which enhances CA release and consequently leads to substantial GSH consumption. While SN38-CA NPs were capable of GSH depletion, this effect was less pronounced without the involvement of FC. The FC-induced Fenton reaction alone was shown to have a mild GSH depleting effect, likely through ROS accumulation and the perturbation of the cellular redox balance. Collectively, SN38-CA@FC NPs demonstrated the strongest capability to deplete GSH levels, which in turn also impacted the expression of intracellular GPX4.

### 2.9 SN38-CA@FC NPs induce intracellular GPX4 depletion

To explore the effect of nanoparticle treatment on GPX4, a crucial enzyme in the defense against oxidative stress, we subjected A549 cells to various formulations and utilized Western blotting to measure intracellular GPX4 levels. As depicted in Fig. 5F, all treatment groups, relative to the control, exhibited a reduction in GPX4 protein levels. The data suggest that GPX4 depletion by FC and SN38-CA acting independently is modest. The SN38-CA@FC group, however, experienced the most significant decline in GPX4, which can be attributed to the substantial release of CA facilitated by the self-amplifying pathway. This pathway, driven by the dual presence of SN38-CA and FC, not only caused extensive redox system disruption but also led to considerable consumption of GPX4. The patterns of GPX4 depletion across the sample groups corresponded with the trends observed in GSH depletion, which aligns with our expectations. This indicates that SN38-CA@FC NPs effectively impair the cellular antioxidant defense, thereby potentiating the ferroptotic cell death pathway.

### 2.10 Biodistribution

The antitumor efficacy of nanoparticles is contingent upon their ability to accumulate within tumors. We thus explored the biodistribution of nanoparticles in xenograft LLC-bearing C57BL/6 mice. Fig. 6A depicts the time-dependent accumulation of SN38-CA@FC NPs, indicated by increased SN38 fluorescence at the tumor site, peaking at 24 h post-administration. Noteworthy accumulation was also observed in the kidneys, liver, and lungs, likely due to the mononuclear phagocyte system’s role in these organs [28]. Compared to the SN38-CA solution, the SN38-CA@FC NPs demonstrated significantly higher tumor accumulation at all measured time points. This can be ascribed to the nanostructure-enhanced permeability and retention effect, facilitating targeted drug delivery to the tumor. These results underscore the SN38-CA@FC NPs’ favorable biodistribution and their efficacy in concentrating within the tumor microenvironment.

**Fig. 6.**
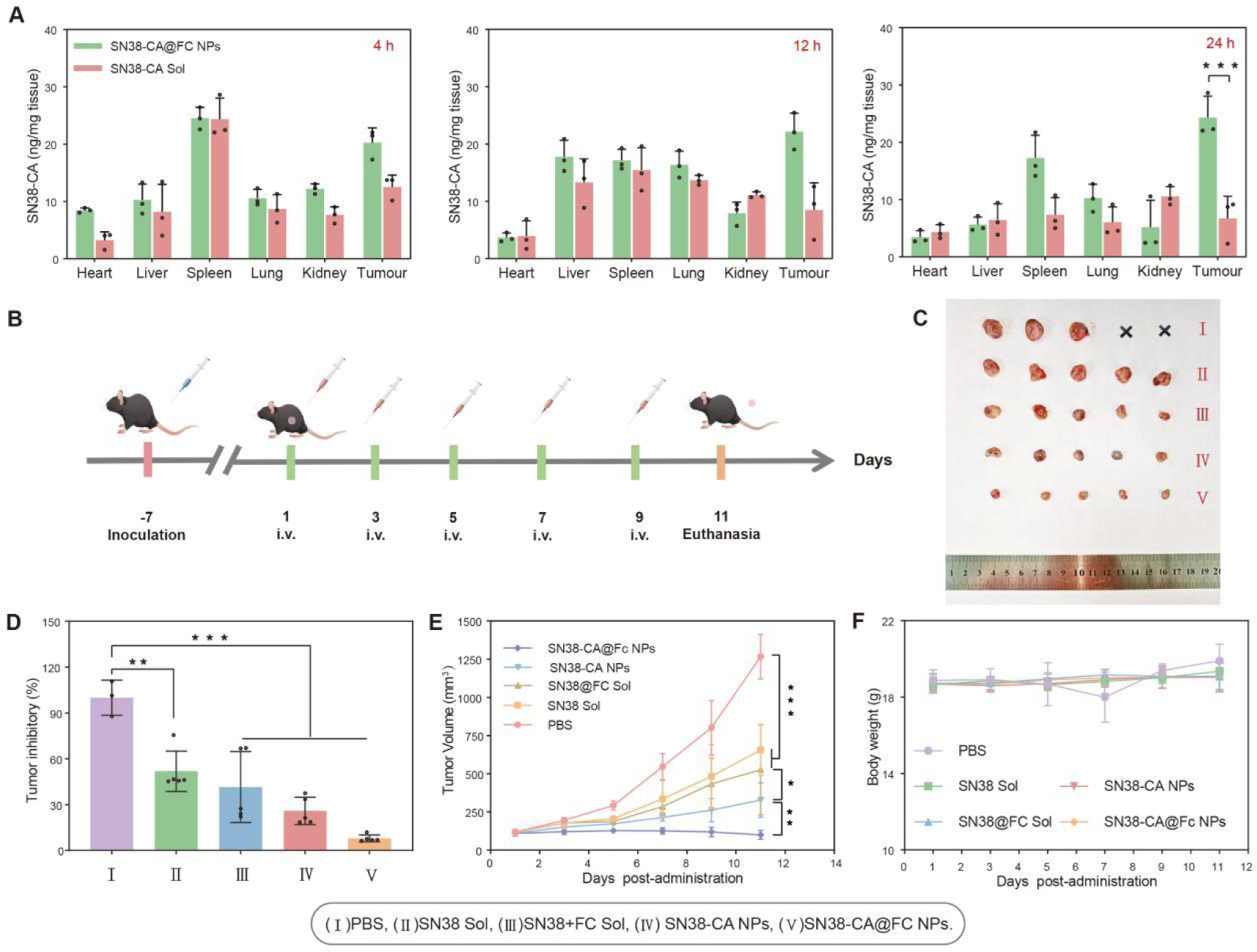
The in vivo performance of the nanoparticles in xenograft LLC-bearing C57BL/6 mice. (A) Biodistribution properties of SN38-CA@FC NPs at 4, 12 and 24 h. (B) Treatment regimen. (C) Photographs of excised tumors. (D) Bar graph depicting tumor inhibition. (E) Tumor volume progression. (F) Body weight fluctuations. ns: no significance, ^∗^*p* < 0.05, ^∗∗^*p* < 0.01, and ^∗∗∗^*p* < 0.001.

### 2.11 In vivo antitumor efficacy

To evaluate the in vivo antitumor efficacy of the SN38-CA@FC NPs, we divided xenograft LLC-bearing C57BL/6 mice into five groups of five. Each group received intravenous injections of PBS, SN38 Sol, SN38+FC Sol, SN38-CA NPs, or SN38-CA@FC NPs, respectively. The treatments were administered on days 1, 3, 5, 7, and 9 (Fig. 6B). Fig. 6C-E show that the PBS group’s tumors grew substantially, with an approximate volume increase of 1250 mm³ by treatment’s end. Moderate tumor growth inhibition was observed with SN38-CA+FC Sol and SN38 Sol, resulting in tumor volumes of about 300-500 mm³. In contrast, both SN38-CA NPs and SN38-CA@FC NPs groups exhibited substantial antitumor activity, with the latter achieving the most pronounced reduction in tumor volume to approximately 100 mm³ at the study’s conclusion. The superior tumor inhibition efficacy was aligned with the trend in tumor volume reduction, with the SN38-CA@FC NPs group showing the greatest inhibitory effect. To assess systemic toxicity, we monitored body weight and hepatorenal function markers (ALT, AST, CRE, and BUN), as depicted in Fig. S8-S11. The nanoparticle treatment did not result in significant alterations in body weight or manifest hepatic or renal injury. Collectively, these findings demonstrate the potent antitumor effects and safety profile of SN38-CA@FC NPs in vivo, which may support their potential clinical application.

## 3. Conclusion

In conclusion, our research has culminated in the creation of a novel self-amplifying, ROS-sensitive SN38 dimeric prodrug nanoparticle, designated as SN38-CA@FC NPs. This innovative formulation synergistically merges chemotherapeutic and ferroptotic mechanisms within a single, efficient NDDS. The synthesized ROS-responsive SN38 dimer prodrug, when combined with FC, yields nanoparticles of uniform size that demonstrate exceptional colloidal stability. The SN38-CA@FC NPs are specifically designed to decompose and release their active components in response to elevated oxidative stress within the tumor microenvironment, while maintaining stability under normal physiological conditions. These nanoparticles effectively augment ROS and lipid peroxide levels, concurrently diminishing GSH and GPX4 concentrations in cancer cells. The culmination of these events is a pronounced antitumor effect both *in vitro* and in animal models, achieved without the compromise of additional systemic toxicity. Thus, our findings present the SN38-CA@FC NPs as a formidable candidate for advanced cancer therapy, with significant potential for clinical translation.

## Acknowledgement

This work was generously provided by the National Natural Science Foundation of China (82104109), Liaoning Provincial Department of Education Program (LJKMZ20221353 and LJKZ0940), Natural Science Foundation of Liaoning Province (2022-BS-158) and the Japan Society for the Promotion of Science (JSPS; 21H01728). The WPI-iCeMS is supported by the World Premier International Research Centre Initiative (WPI), MEXT, Japan.

