## Supplementary Information for "Self-amplifying ROS-sensitive SN38 Dimeric Prodrug nanoparticles for Combined Chemotherapy and Ferroptosis in Cancer Treatment"

**Materials and Methods**

**Materials**

SN38, 2-Mercaptoethylalcohol, 4-Nitrophenyl chloroformate were purchased from Energy Chemical Co., ltd (Shanghai, China). CA, Poloxamer F127, and dithio-dinitrobenzoic acid (DTNB) were purchased from Heowns Biochemical Technology Co., ltd (Tianjin, China). Penicillin, streptomycin, DMEM cell culture medium, fetal bovine serum (FBS), phosphate buffered solution (PBS), Trypsin-EDTA, reactive oxygen species assay kit (DCFH-DA), Hoechst 33342, and the reagents applied in Western blot assay were obtained from Dalian Meilun Biotech Co., ltd (China). 3-(4,5-dimethylthiazol-2-yl)-2,5-diphenyl tetrazolium bromide (MTT) were purchased from Biosharp Biotechnology Co., ltd (Anhui, China). Deferoxamine (DFO) was purchased from Bidepharm Technology Co., ltd (Shanghai, China). The antibodies C11 BODIPY ^581/591^ were purchased from Shanghai Maobin Medical Technology Co., ltd (Shanghai, China). GSH and GSSG Assay Kit was purchased from Biyuntian Biotechnology Co., ltd (Shanghai, China). Enzyme-Linked Immunosorbnent Assay (ELISA) kit (ALT, AST, CRE and BUN) was purchased from Nanjing Jiancheng Bioengineering Institute. All the vessels for cell culture were provided by Dalian Meilun Biotech Co., ltd (China) and any other solvents and chemicals were of analytical or HPLC grade.

**Cells and animals**

A549 and LLC cells were cultured in DMEM medium supplemented with 10% FBS at 37℃ in 5% CO_2_ atmosphere. C57BL/6 mice (18-22 g, female) were purchased from Liaoning Changsheng Biotechnology Co., ltd (Benxi, China). All animal experiments were approved by the Institutional Animal Care and Use Committee of Shenyang Pharmaceutical University. For developing the xenograft LLC tumor model, 2 × 10^7^ cells/1 ml PBS were subcutaneously injected into the right mammary fat pads of the mice.

**Synthesis of the prodrug**

The synthesis route of SN38-CA is as shown in Fig. 2. Firstly, CA (1 g, 7.57 mmol) and 2-mercaptoethylalcohol (8.85 g, 113.27 mmol) were co-dissolved in ethyl acetate under the condition of ice bath, and zirconium chloride (17.65 mg, 0.076 mmol) was added as catalyst, stirring at room temperature for 30 min. The reaction solution was washed with 10% (v/v) citric acid solution and saturated sodium chloride solution, respectively, the organic layer was collected, and the crude product was obtained by Rotary evaporator. The Midbody-1 was purified by silica gel column chromatography (petroleum ether (PE): ethyl acetate (EA) = 1.5:1, v/v). Subsequently, the Midbody-1 (200 mg, 0.74 mmol), 4-Nitrophenyl chloroformate (650 mg, 3.22 mmol) and (Triethylamine) TEA (150 mg, 1.48 mmol) were dissolved in dichloromethane (DCM) and reacted at room temperature for 1 h. The reaction solution was washed with 10% (v/v) NaHCO_3_ solution, 10% (v/v) citric acid solution and saturated sodium chloride solution, respectively, the organic layer was collected, and the crude product was obtained by Rotary evaporator. The Mmidbody-2 was purified by silica gel column chromatography (petroleum ether (PE): ethyl acetate (EA) = 8:1, v/v). Finally, Midbody-2 (25 mg, 0.042 mmol), SN38 (50 mg, 0.13 mmol) and TEA (21 mg, 0.21 mmol) were dissolved in N, N'-dimethylformamide (DMF) and reacted at room temperature for 24 h. The reaction solution was washed with 10% (v/v) Na_2_CO_3_ solution, 10% (v/v) citric acid solution and saturated sodium chloride solution, respectively, the organic layer was collected, and the crude product was obtained by Rotary evaporator. SN38-CA was purified using Preparative liquid chromatograph (acetonitrile: ethanol = 65:35, v/v) to obtain SN38-CA.

**Preparation of nanoparticles**

The method of preparing nanoparticles is the classical one-step nanoprecipitation method.In brief, a solution containing 1 mg of SN38-CA and 0.25 mg of Poloxamer F127 was dissolved in 300 μL of acetone. The drug-containing solution was then slowly added dropwise to a 2 mL aqueous solution under vigorous stirring for 15 min. Subsequently, the organic solvent was removed via rotary evaporation to obtain the SN38-CA nanoparticles. The SN38-CA@FC NPs were prepared using the same procedure, with the addition of FC (1 mg) to the drug-containing solution. The particle size, polydispersity index (PDI) and zeta potential of prodrug nanoparticles were measured using Malvern Nano Zetasizer. The morphology of nanoparticles was examined utilizing a Tecnai G220 Transmission Electron Microscope (TEM). The existence of Tyndall effect was detected by red laser irradiation.

**In vitro Fenton reaction detection**

Crystal violet spectrophotometry was used to detect the occurrence of Fenton reaction in vitro. The principle is that crystal violet can react electrophilic with hydroxyl radicals produced by Fenton reaction, and the absorbance decreases. The experiment consisted of four groups: crystal violet, crystal violet+H_2_O_2,_ and crystal violet+H_2_O_2_+FC. After incubation at 37℃ for 30 minutes, the absorbance of each group at 580 nm was detected.

**Colloidal stability of prodrug nanoparticles**

To investigate the colloidal stability of SN38-CA NPs and SN38-CA@FC NPs under physiological conditions, we cultured the samples in PBS supplemented with 10% (v/v) FBS at pH 7.4 and incubated them at 37°C with continuous shaking. The samples were collected at time points of 0, 0.5, 1, 2, 4, 6, 8 and 12 h respectively for particle size analysis.

**Study on assembly Mechanisms of Prodrug nanoparticles**

The masking intermolecular force reagent with 900 μL concentration of 1 M was added to 100 μL nanoparticle solution, and the particle size and PDI changes were measured by DLS at time points: 0, 0.5, 1, 2, 4, 6 and 8 h.

**Evaluation of in vitro drug release of prodrug nanoparticles**

To investigate the ROS responsiveness of SN38-CA NPs and SN38-CA@FC NPs, we used a medium comprising PBS-ethanol solution (70: 30, v/v) with or without H_2_O_2_ as stimuli (10 mM). Briefly, SN38-CA NPs and SN38-CA@FC NPs (0.25 mg/mL, SN38 equivalent,2 mL) were placed in the release medium (8 mL) at 37°C under shaking. At selected time intervals (0, 1, 2, 4, 8, 12 and 24 h),0.6 mL samples were collected (n = 3) and 0.6 mL of blank release medium were subsequently replenished. The concentration of released SN38 was determined with high performance liquid chromatography at a wavelength of 365 nm.

**In vitro cytotoxicity assay**

The in vitro cytotoxicity of A549 and LLC cells was assessed using a standard MTT assay. Briefly, 2.5 × 10^4^ cells per well were seeded into 96-well plates and incubated overnight before adding SN38 Sol, SN38+Fc Sol, SN38-CA NPs or SN38-CA@FC NPs at varying concentrations for a 24 h incubation period. Afterward, 10 μL of MTT solution (5 mg/mL) was added to each well and incubated at 37℃ for 4 h . Subsequently, the medium containing MTT solution was aspirated and replaced with 200 μL of dimethyl sulfoxide (DMSO) per well to dissolve formazan crystals for a duration of 10 min. Consequently, the absorbance at 490 nm was measured using a multifunctional micropore detection board analysis system. The IC_50_ value was calculated utilizing GraphPad 8.0 software.

**Intracellular ROS assay**

The intracellular level of ROS was assessed using DCFH-DA, which can be oxidized in the presence of ROS to generate a fluorescent compound called dichlorofluorescin (DCF). Briefly, A549 cells were seeded at a density of 2.5 × 10^5^ cells per well in DMEM medium supplemented with 10% (v/v) FBS and incubated overnight. Subsequently, the cells were treated with PBS, SN38+Fc Sol, SN38-CA@Fc NPs + DFO (100 µM), SN38-CA NPs and SN38-CA@FC NPs at the SN38-equivalent concentration of 1 µM, respectively. After incubating for 4 h, the cells were washed with PBS and stained with DCFH-DA (20 µM) for 30 min. Finally, the cells were washed with PBS, covered with antifade mounting medium and studied by Confocal Laser Scanning Microscopy (CLSM).

**Intracellular LPO assay**

The intracellular LPOs level was monitored using C11-BODIPY^581/591^ as an indicator and analyzed by CLSM. Briefly, A549 cells were seeded into 6-well plates at a density of 2.5 × 10^5^ cells in DMEM medium with 10% FBS and incubated overnight. Subsequently, the cells were treated with PBS, SN38+Fc Sol, SN38-CA@Fc NPs + DFO (100 µM), SN38-CA NPs, and SN38-CA@FC NPs at a concentration equivalent to 1 µM of SN38. After a 4 h incubation period, the cells were rinsed with PBS and subsequently stained with C11-BODIPY^581/591^ (20 µM) for 30 min. Finally, the cells were washed again with PBS, covered with antifade mounting medium, and examined using CLSM.

**Extracellular GSH assay**

The ability of extracellular CA to consume GSH was examined by DTNB. The principle is that DTNB can react with the sulfhydryl group (-SH) of GSH to produce a colored substance, TNB, which has a strong absorbance at 412 nm. When GSH is oxidized to GSSH, it cannot react. The control group(150 µg/mL GSH sol+PBS) and CA group(150 µg/mL GSH sol+500 µg/mL CA sol) were incubated at 37ºC. The samples were taken out at 0, 0.5, 1 and 2 h, and incubated at 37ºC for 5 min after adding DTNB solution, and the absorbance at 412 nm was detected.

**Intracellular** **GSH assay**

The intracellular GSH content was assessed using a GSH kit. Briefly, A549 cells were seeded into 6-well plates at a density of 2.5 × 10^5^ cells in DMEM medium supplemented with 10% (v/v) FBS and incubated overnight. Subsequently, the cells were treated with PBS, SN38+Fc Sol, SN38-CA@Fc NPs + DFO (100 µM), SN38-CA NPs, and SN38-CA@FC NPs at an equivalent concentration of 1 µM of SN38, respectively. After incubation for 4 h, the cells were washed with PBS, centrifuged, and the supernatant was discarded. The cells precipitation was then mixed thoroughly with three times the volume of protein removal agent M solution. Finally, three times the volume of protein removal agent M solution was added to the cell pellet and vortexed completely. The cell samples were rapidly frozen and thawed twice using liquid nitrogen and a 37 ºC water bath. Subsequently, the samples were placed at 4 ºC for 5 min and centrifuged at 4 ºC for 10 min at a speed of 10000 rpm. The resulting supernatant was utilized to determine the total glutathione concentration. Following the removal of GSH, the content of GSSG was determined, and the content of GSH was equal to the total glutathione content minus the GSSG content.

**Intracellular GPX4 assay**

The change in GPX4 protein expression in A549 cells was detected using Western blot analysis. Briefly, A549 cells were seeded into 6-well plates at a density of 2.5 × 10^5^ cells in DMEM medium supplemented with 10% (v/v) FBS and incubated overnight. Subsequently, the cells were treated with PBS, SN38+Fc Sol, SN38-CA@Fc NPs +DFO (100 µM), SN38-CA NPs and SN38-CA@FC NPs at the SN38-equivalent concentration of 1 µM, respectively. After incubating for 12 h, the cells were lysed and the protein concentration was determined using the BCA Protein Assay Kit. The protein was then separated by 12% electrophoresis, transferred onto nitrocellulose filter membranes (Millipore, U.S.A.), and subsequently blocked with 4% bovine serum albumin for 2 h to prevent nonspecific binding. Subsequently, the membranes were incubated with diluted primary antibodies GPX4 (1:1000) at 4 ºC for 12 h. After thorough washings, the membranes were then incubated for 2 h with a secondary antibody, goat anti-rabbit immunoglobulin G, and GAPDH (1:5000), following standard protocols. Finally, the expression of GPX4 was quantified using the GEA Mersham Imager 600 and enhanced chemiluminescence working solution.

**Biodistribution study**

The in vivo biodistribution of SN38-CA@FC NPs was investigated in C57BL/6 mice bearing xenograft LLC tumors. After tumor volume reached 200 mm^3^, the tumor-bearing mice were randomly divided into two groups (n = 3) and intravenously injected with 200 μL of SN38-CA Sol and SN38-CA@FC NPs at SN38-CA at a dose of 10 mg/ kg, respectively. Major organs including the heart, liver, spleen, lungs and kidneys, as well as tumors were collected at 4 h, 12 h and 24 h post-injection. Major organs were washed with PBS, dried and subsequently weighed. Then, the major organs were homogenized by tissue homogenizer, and the content of SN38-CA in the homogenizer was detected by enzyme label.

**In vivo antitumor efficacy study**

The in vivo antitumor efficacy of SN38-CA@FC NPs was investigated in the xenograft LLC bearing C57BL/6 mice. After the tumor volume reached 100 mm^3^, the tumor-bearing mice were randomly divided into five groups (n = 5) and intravenously injected with different treatments: (1) PBS, (2) SN38 Sol, (3) SN38+FC Sol, (4) SN38-CA NPs, and (5) SN38-CA@FC NPs at a dose of 5 mg/kg on days 1, 3, 5, 7, and 9. The tumor volumes and body weights were measured bi-daily. The tumor volume was calculated using the formula: *V* = *L* × *W* × *W*/2 (where L represents the longest dimension and W represents the shortest dimension). On day 11, the mice were euthanized for further examination. Liver function (Alaninetransaminase,ALT; Aspartate aminotransferase, AST) and kidney function (Blood urea nitrogen, BUN; Complete response, CR) were measured by the hepatorenal function kit with blood taken from the eyeball.

**Statistical analysis**

Data are presented as mean ± SD. Significant difference was analyzed by one-way ANOVA. The following p-values were considered to be significant: ∗*p* < 0.05, ∗∗*p* < 0.01, and ∗∗∗*p* < 0.001.

**Supplementary Figures**

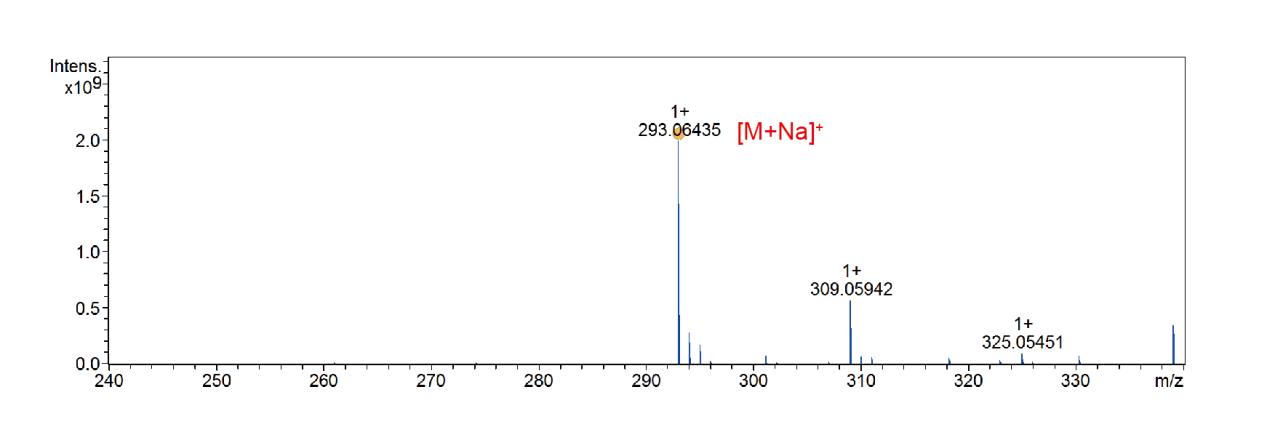

**Fig. S1:** MS spectrum of midbody 1, [M+Na]^+^=293.06435.

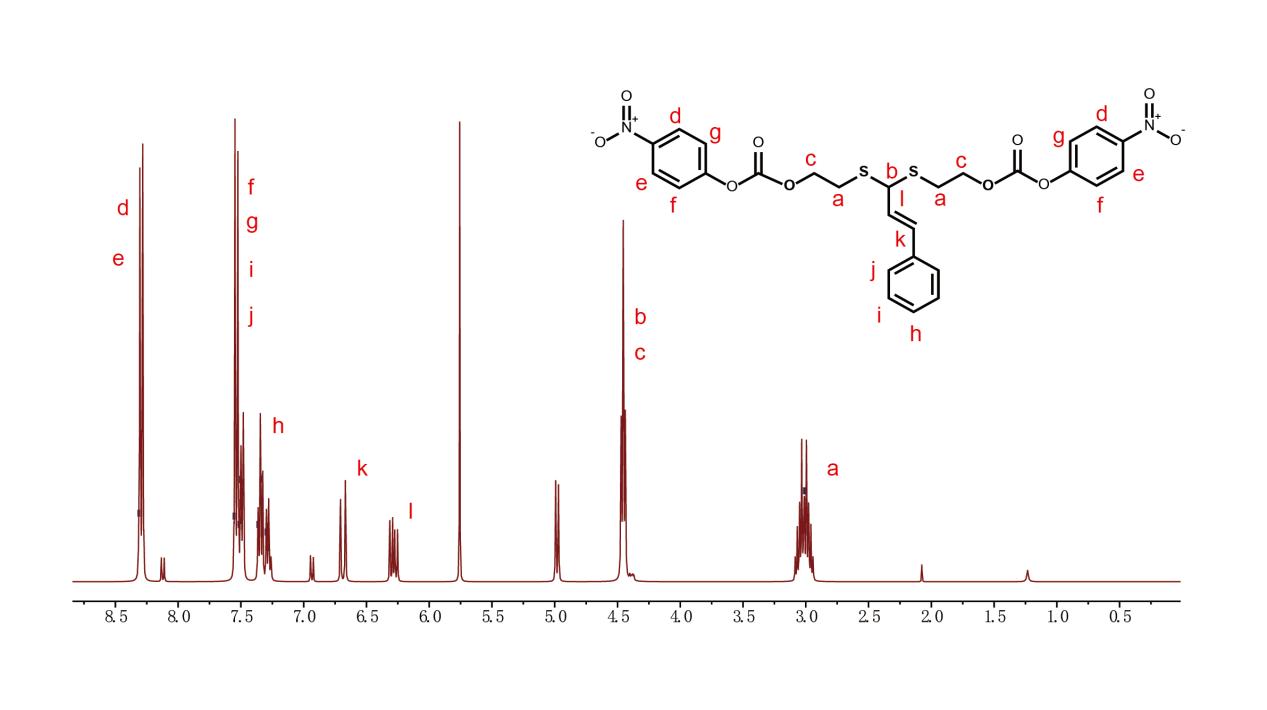

**Fig. S2.** ^1^H NMR spectrum of midbody 2 in DMSO-*d6*.

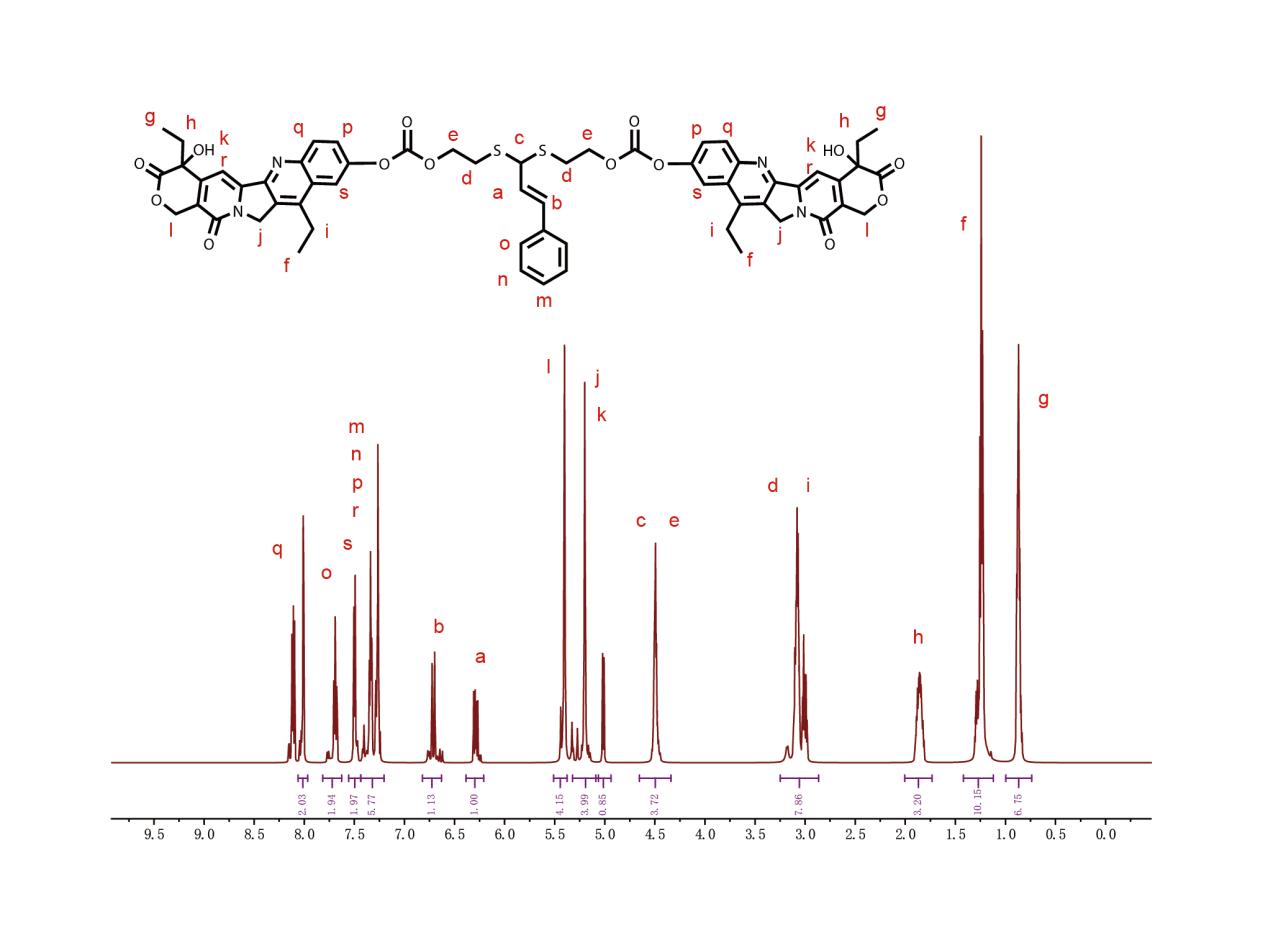

**Fig. S3.** ^1^H NMR spectrum of SN38-CA in DMSO-*d6*.

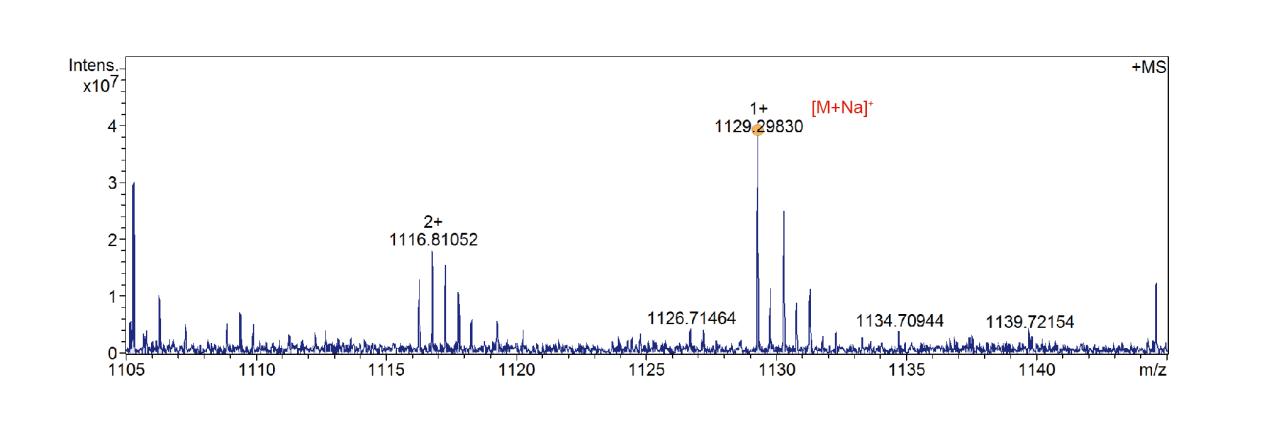

**Fig. S4:** MS spectrum of SN38-CA, [M+Na]^+^=1129.29830.

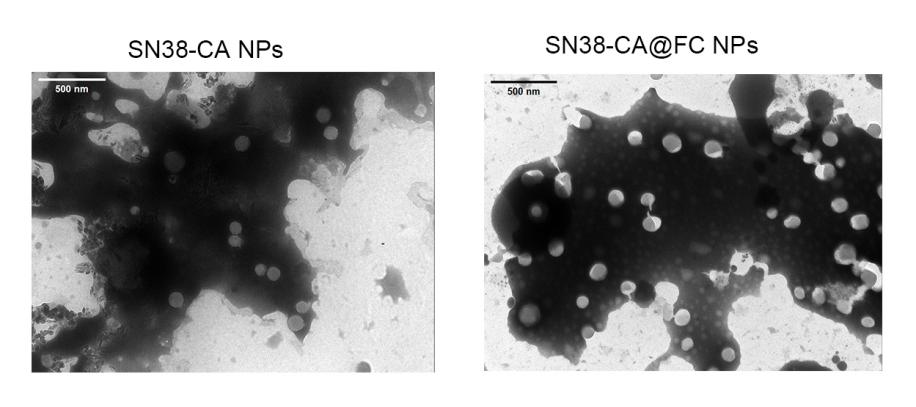

**Fig. S5:** The morphology of SN38-CA NPs and SN38-CA@FC NPs was photographed by TEM.

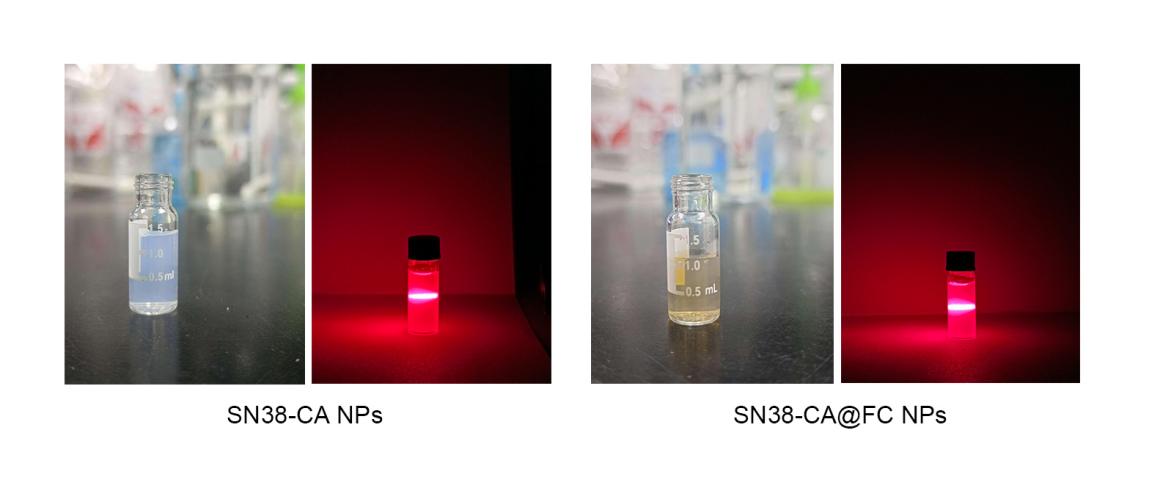

**Fig. S6:** The appearance and the Dundahl effect picture of SN38-CA NPs and SN38-CA@FC NPs.

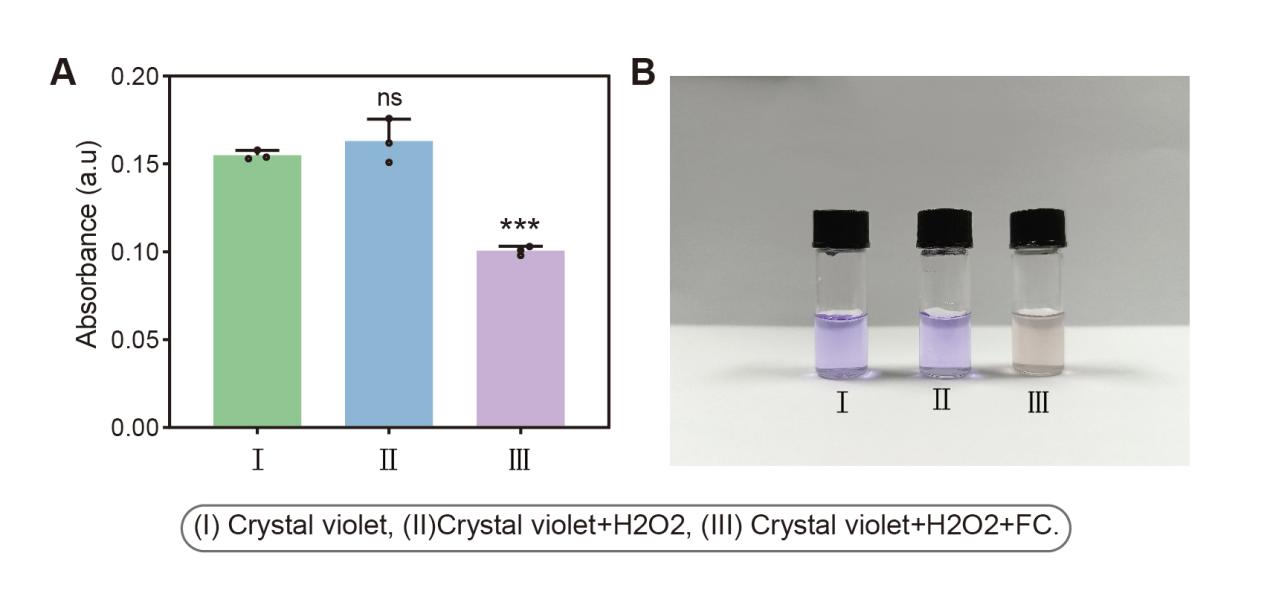

**Fig. S7:** In vitro Fenton reaction detection. (A) Absorbance of each group at 480nm after incubation at 37℃ for 30min. (B) Appearance of each group at 480nm after incubation at 37℃ for 30min.

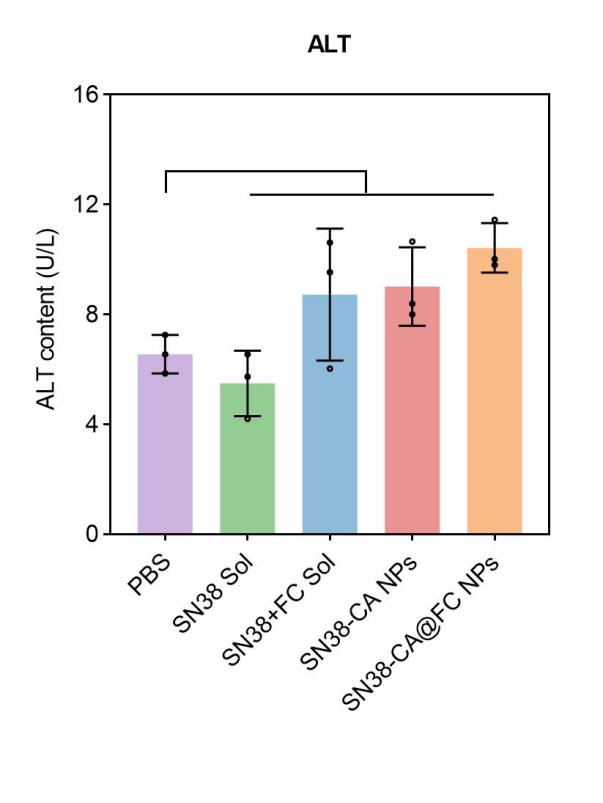

**Fig. S8**. ALT content of xenograft LLC-bearing C57BL/6 mice treated with different formulations.

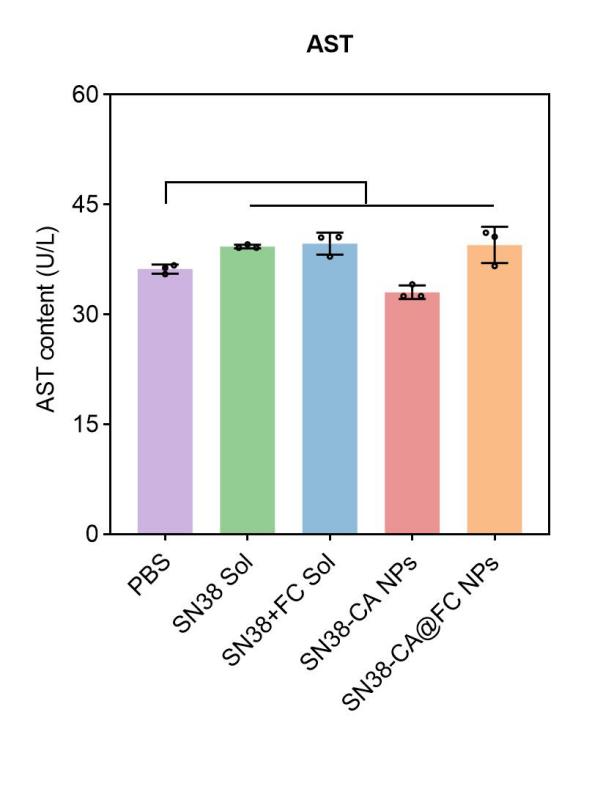

**Fig. S9**. AST content of xenograft LLC-bearing C57BL/6 mice treated with different formulations.

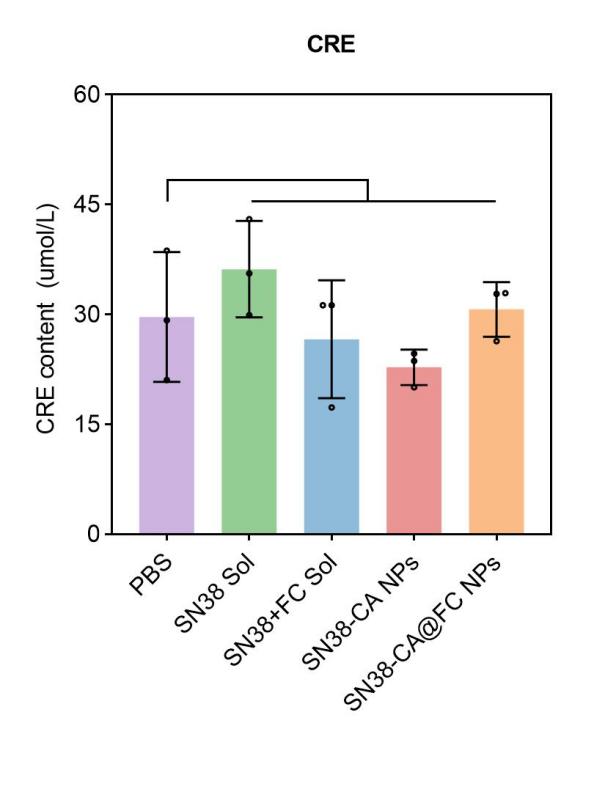

**Fig. S10**. CRE content of xenograft LLC-bearing C57BL/6 mice treated with different formulations.

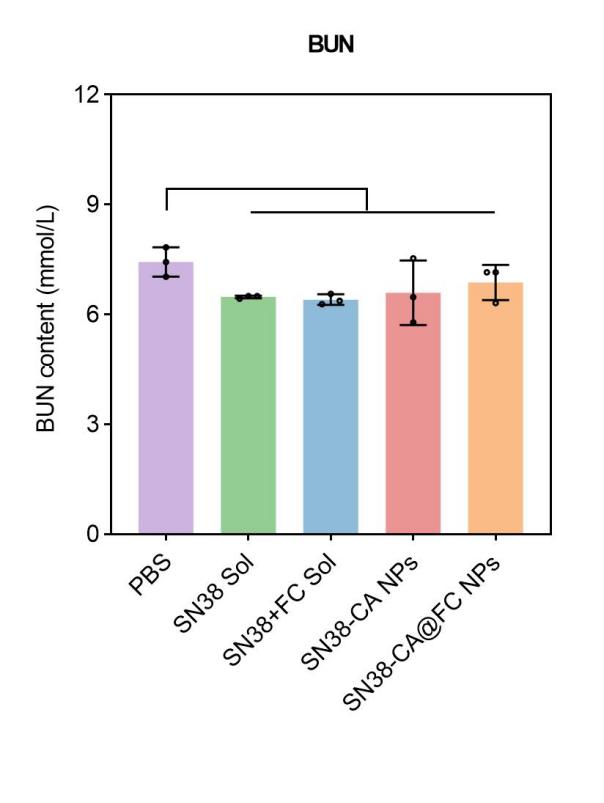

**Fig. S11**. BUN content of xenograft LLC-bearing C57BL/6 mice treated with different formulations.

**Tab. S1.** IC_50_ value of different formulations against A549 cells.

| Formulations | IC50 (nM) |
| --- | --- |
| SN38 Sol | 192.50 |
| SN38@FC Sol | 62.43 |
| SN38-CA NPs | 37.28 |
| SN38-CA@FC NPs | 11.93 |

**Tab. S2.** IC_50_ value of different formulations against LLC cells.

| Formulations | IC50 (nM) |
| --- | --- |
| SN38 Sol | 181.30 |
| SN38@FC Sol | 147.70 |
| SN38-CA NPs | 107.50 |
| SN38-CA@FC NPs | 77.68 |
